# Genome sequencing and comparative analysis of *Ficus benghalensis* and *Ficus religiosa* trees reveal evolutionary mechanisms of longevity

**DOI:** 10.1101/2021.10.14.464369

**Authors:** Abhisek Chakraborty, Shruti Mahajan, Manohar Singh Bisht, Vineet K. Sharma

## Abstract

*Ficus benghalensis* (Indian banyan tree) and *Ficus religiosa* (Peepal) trees are well-known for their long lifespan, traditional significance, and medicinal properties. Therefore, to understand the genomic and evolutionary aspect of these characteristics the whole genomes of these two *Ficus* species were sequenced using 10x Genomics and Oxford Nanopore sequencing platforms. The draft genome assemblies of *F. benghalensis* (392.89 Mbp genome containing 25,016 high-confidence coding genes), and *F. religiosa* (332.97 Mbp genome containing 23,929 high-confidence coding genes) were constructed. We also established the genome-wide phylogenetic position of the two *Ficus* trees with respect to 46 other Angiosperm plant species and studied the comparative population demographic history of these two species to show a population bottleneck event ~0.8 Mya for both the species. We also identified 7,468 orthogroups across 16 phylogenetically closer Eudicot plant species including *F. benghalensis* and *F. religiosa*. Comparative evolutionary analyses using these orthogroups, gene family expansion/contraction analysis, and gene duplication analysis showed adaptive evolution in genes involved in cellular pathways and mechanisms that are central to plant growth and development and provide genomic insights into longevity and ecological significance of these large woody trees.

## INTRODUCTION

*Ficus* is one of the most diverse genus that originated around 80-90 million years ago, and consists of ~800 species^1,2^. *Ficus* plants belong to the plant family Moraceae, also known as fig family consisting of over 1,100 species^3^. Plants from this genus are also known as fig plants for the presence of their characteristic inflorescence known as syconium or fig that helps in pollination of fig plants with the help of pollinators such as fig wasps^1^. *Ficus* plants are abundant in a wide range of tropical and subtropical ecosystem. Fig plants possess several characteristics such as smaller seed size, higher growth rate, higher fecundity etc. that make them evolutionarily flexible to grow in diverse environmental habitats^2^. Species from the *Ficus* genus occupy diverse ecological niches, and different life forms such as free-living, epiphytes, semi-epiphytes, lithophytes, or rheophytes^4^. Fig plants are considered as keystone species in a tropical ecosystem as they sustain a large number of animal species with fig fruits that are produced throughout the year^5^. Large *Ficus* plants (including *F. benghalensis* and *F. religiosa*) are crucial components of tropical ecosystem as they can be considered as forest restoration agents due to their larger canopy areas that favor plant species dispersed by frugivores as well as other species^6^. Fig plants, and their pollinator species such as fig wasps coevolved in course of time to show obligate mutualism^7^. For pollination purpose, fig plants interact with their wasp pollinators by attracting them with volatile organic compounds (VOCs)^7^.

*F. benghalensis*, also known as Indian banyan tree, is the National tree of India, and is native to tropical, subtropical regions in South Asia^8^. It is also known for its traditional religious, mythological significance in Hinduism and Buddhism, and their wide-spread branches signify eternal life^8^. *F. benghalensis* (a strangler fig) is a large woody plant with around 30 meters height, and typically have a long life spanning several centuries^9^. The largest banyan tree (Thimmamma Marrimanu) is located in Southern India, and is 550 years old^9^. It starts growing as an epiphyte in early days, and in course of time it disperses thick plentiful aerial roots from the spreading branches to cover several hundred meters and grows independently^10^. The extracts from bark, leaf and fruit of this plant possess various bioactive compounds e.g., flavonoid, saponin, sterol, anthocyanidin derivatives etc. that confer a large number of pharmacological activities e.g., anti-inflammatory, anti-helminthic, anti-oxidant, anti-microbial, anti-tumor, antipyretic, immunomodulatory, and wound healing properties^8^.

*F. religiosa*, also known as ‘sacred fig’, ‘Bodhi tree’, ‘Peepal’, or ‘Ashwattha’, is a large hemi epiphyte, deciduous tree with long average lifespan of 900-1,500 years^11,12^. It is also a well-known tree for its mythological and religious importance in Buddhism and Hinduism, and is known as the oldest tree in Hindu mythology^12,13^. Similar to *F. benghalensis*, the *F. religiosa* is also a large perennial tree with up to 20 meters height, is epiphytic in young stages and the adult tree have widespread branches but without the aerial roots^11,14^. Different parts of this plant such as root bark, stem bark, fruits, and seeds contain essential compounds e.g., steroids, alkaloids, flavonoids, phenols, tannins, essential amino acids that have a wide range of pharmacological applications^11^. Phytochemical analysis of the barks of *F. religiosa* showed the presence of bioactive compounds such as flavonoids, glycosides, terpenoids, tannins, saponins, wax, and others^15^ that can also be used as potential therapeutic agents^16^. These components are responsible for conferring anti-oxidant, anti-bacterial, hypoglycemic, hypolipidemic, wound-healing, antihelminthic, immunomodulatory, anti-diabetic, anti-convulsant, and anti-ulcer activities to *F. religiosa* plant^11,15^.

*F. benghalensis* and *F. religiosa* genomes are diploid (2n=26) and the estimated genome size is 686 Mbp for both the species according to Plant DNA C-values Database^17^. A few *Ficus* species such as *F. carica, F. hispida, F. microcarpa* and *F. erecta* have been sequenced^7,18,19^; however these two significant *Ficus* genomes are yet not sequenced. In this study, we carried out the first genome sequencing and analysis of *F. benghalensis* and *F. religiosa* using 10x Genomics linked reads and Oxford Nanopore long reads to get evolutionary insights into the mechanisms related to medicinal properties and long lifespan. We also performed transcriptome sequencing of these species that helped in comprehensive genome annotation. The phylogenetic position of the two fig species was resolved using other available Eudicot species, and comparative evolutionary analysis with selected Angiosperm plant species revealed that genes required for plant growth, development, and stress tolerance mechanisms are highly evolved in these species, which are perhaps responsible for the longevity of these two plant species.

## RESULTS

### Genome and Transcriptome Sequencing

Genome sizes of *F. benghalensis* and *F. religiosa* were estimated as 431.53 Mbp and 413.14 Mbp, respectively, using SGA-preqc^20^. Supernova^21^ v2.1.1 also predicted the genome sizes as 420.13 Mbp and 388.12 Mbp for *F. benghalensis* and *F. religiosa*, respectively. A total of 49 Gb and 9 Gb of genomic data was generated from *F. benghalensis* leaf tissue using 10x Genomics (Chromium) and Oxford Nanopore technology, respectively. Genomic data corresponded to 113.6X coverage of 10x Genomics linked read data, and 20.9X coverage of Nanopore long read data based on predicted genome size using SGA-preqc. Further, a total of 7.4 Gb of RNA-Seq data was also generated for *F. benghalensis* plant species (**Supplementary Tables 1-2**). For *F. religiosa* plant species, 10x Genomics and Oxford Nanopore sequencing technologies were used to generate 48.7 Gb (117.8X) and 7.2 Gb (17.4X) of genomic data, respectively. Further, a total of 10.8 Gb of RNA-Seq data was also generated (**Supplementary Tables 1-2**).

### Genome assembly

The heterozygosity content for *F. benghalensis* was estimated to be 1.78% (**Supplementary Figure 1a**). The final genome assembly of *F. benghalensis* (assembly v3) had a total size of 392.89 Mbp with 4,822 contigs, N50 value of 486.9 Kbp, longest contig size of 5.4 Mbp, and a GC-content of 34.54% (**Supplementary Table 3**). Final genome assembly size was close to the estimated genome size of 381.9 Mbp predicted using GenomeScope^22^ (**Supplementary Figure 1a**). *F. benghalensis* genome assembly had 96.4% complete BUSCOs and 1.7% of fragmented BUSCOs (**Supplementary Figure 1c, Supplementary Table 4**). 98.39% of the barcode-filtered linked-reads and 93.11% adapter-processed Nanopore reads could be mapped on the final genome assembly.

The heterozygosity content for *F. religiosa* genome was estimated to be 1.62% (**Supplementary Figure 1b**). *F. religiosa* final genome assembly had a total size of 332.97 Mbp consisted of 6,087 contigs (largest contig of 4.51 Mbp), an N50 value of 553.4 Kbp, and a GC-content of 34.31% (**Supplementary Table 3**). This final *F. religiosa* genome assembly had 95.6% complete BUSCOs, and 1.1% of fragmented BUSCOs (**Supplementary Figure 1c, Supplementary Table 4**). Further, 98.14% of the barcode-filtered 10x Genomics linked reads and 93.99% of the adapter-processed Nanopore long reads could be mapped to the final genome assembly of *F. religiosa*. Final assembled genome size of *F. religiosa* was close to the predicted genome size of 337.7 Mbp using GenomeScope^22^ (**Supplementary Figure 1b**).

Sequence variation analysis in these final genome assemblies using barcode-filtered 10x Genomics reads identified 1,772,802 nucleotide positions (0.45%) as variant sites in *F. benghalensis*, and 1,466,212 variant sites (0.44%) in *F. religiosa*. These variant sites were located in 3,254 and 3,859 contigs (after scaffolding) in *F. benghalensis* and *F. religiosa*, respectively.

### Genome annotation

A total of 2,159 repeat family sequences were identified in *F. benghalnesis* genome assembly, which were clustered to obtain 1,879 repeat families that were used to repeat-mask the *F. benghalensis* genome. Using this *de novo* repeat library, 51.36% of the genome assembly was masked as repetitive regions using RepeatMasker. Among these, 47.08% was identified as interspersed repeats, which was further divided into 32.88% unclassified, 11.81% retroelements (including 2.70% Ty1/Copia, and 8.28% Gypsy/DIRS1 elements), and 2.39% DNA transposons (**Supplementary Figure 1d, Supplementary Table 5**). 7.01% of the genome was also detected as tandem repeats using TRF^23^ v4.09. Thus, ~54% of the *F. benghalensis* genome was predicted to consist of simple and interspersed repeat regions. Additionally, 1,028 rRNAs, 679 tRNAs (decoding standard amino acids), and 195 miRNAs were identified in *F. benghalensis* genome.

Similarly, 2,002 *de novo* repeat family sequences were identified in *F. religiosa* final genome assembly, which were clustered to obtain 1,735 repeat sequences that were used as *de novo* repeat library. Repeat-masking of *F. religiosa* genome assembly was performed using this library. RepeatMasker analysis masked 45.88% of the *F. religiosa* genome assembly as repetitive regions (41.68% interspersed repeats). Additionally, Tandem Repeat Finder (TRF) also predicted 8.84% of the genome sequence as simple repeats. Among the interspersed repeats, 6.70% was retroelements (1.55% Ty1/Copia, 4.51% Gypsy/DIRS1 elements), 1.64% was DNA transposons, and 33.35% was unclassified (**Supplementary Figure 1d, Supplementary Table 6**). Therefore, ~51% of the *F. religiosa* genome assembly consisted of repetitive regions (including both simple and interspersed repeats). Also, 668 tRNAs (decoding standard amino acids), 1,135 rRNAs, and 185 miRNAs were detected in the final *F. religiosa* genome assembly.

Prior to gene set construction, quality-filtered RNA-Seq data for *F. benghalensis* plant were *de novo* assembled, which resulted in a total of 10,156 transcripts (used as a set of empirical evidence in MAKER pipeline^24^). A total of 29,524 gene models were identified from MAKER genome annotation pipeline. Among these, 27,674 genes (93.73%) having an AED value of <0.5 were extracted. Among these 27,674 genes, 25,016 were filtered out based on the length-based criteria of ≥300 bp and were retained in the high-confidence gene set of *F. benghalensis*. In case of *F. religiosa* species, quality-filtered RNA-Seq data was used for *de novo* transcriptome assembly resulting in 74,514 assembled transcripts, which were used for evidence-based alignments in MAKER genome annotation pipeline. The pipeline predicted a total of 27,544 coding gene sequences, among which, 26,126 genes (94.85%) had AED values <0.5. After extracting these genes, and further length-based filtering (≥300 bp), a total of 23,929 coding genes were retained to construct the high-confidence gene set for *F. religiosa* genome.

In the resultant coding gene set of *F. benghalensis*, 86.3% genes were identified as complete BUSCOs and 8.1% genes were identified as fragmented BUSCOs. Out of 25,016 high-confidence genes, 24,909 genes, 20,947 genes, and 20,541 genes could be mapped against NCBI-nr, Swiss-Prot, and Pfam-A databases, respectively, and 90 genes (0.36%) could not find a match in any of these databases. In case of *F. religiosa*, the resultant coding gene set had 90.1% complete BUSCOs, and 4.2% of fragmented BUSCOs. Among the 23,929 high-confidence coding genes, 23,638 genes, 19,850 genes, and 19,437 genes found matches against NCBI-nr, Swiss-Prot, and Pfam-A databases, respectively. Overall, a total of 262 genes (1.09%) could not be annotated using any of these databases.

### Synteny analysis

Intra-species collinear blocks were identified using MCScanX^25^, which showed 23.72% collinearity (5,933 out of 25,016 genes) among *F. benghalensis* coding genes and 18.65% collinearity (4,463 out of 23,929 genes) among *F. religiosa* coding genes. Further, 48.44% dispersed, 4.37% proximal, 12.93% tandem, and 23.72% segmental duplicated genes, and 10.54% singleton genes were detected in *F. benghalensis* coding gene set. *F. religiosa* contained 53.21% dispersed, 3.63% proximal, 12.99% tandem, 18.65% segmental duplicated genes, and 11.52% singleton coding genes.

503 inter-species syntenic blocks were identified, which involved 58.33% of the total coding genes predicted for *F. benghalensis* and *F. religiosa*, including 14,688 *F. benghalensis* syntelogs (58.71% of the predicted genes) and 13,861 *F. religiosa* syntelogs (57.93% of the predicted genes).

### Demographic history and Phylogenetic position of *Ficus* species

Effective population size (N_e_) estimation for the two *Ficus* species using PSMC^26^ analysis showed that both the species suffered one major population bottleneck in a similar time period - around 0.8 Million years ago **(Figure 1a)** in the Pleistocene epoch of the Geological time scale, which is similar to the population demographic history of other tropical forest plants^27^.

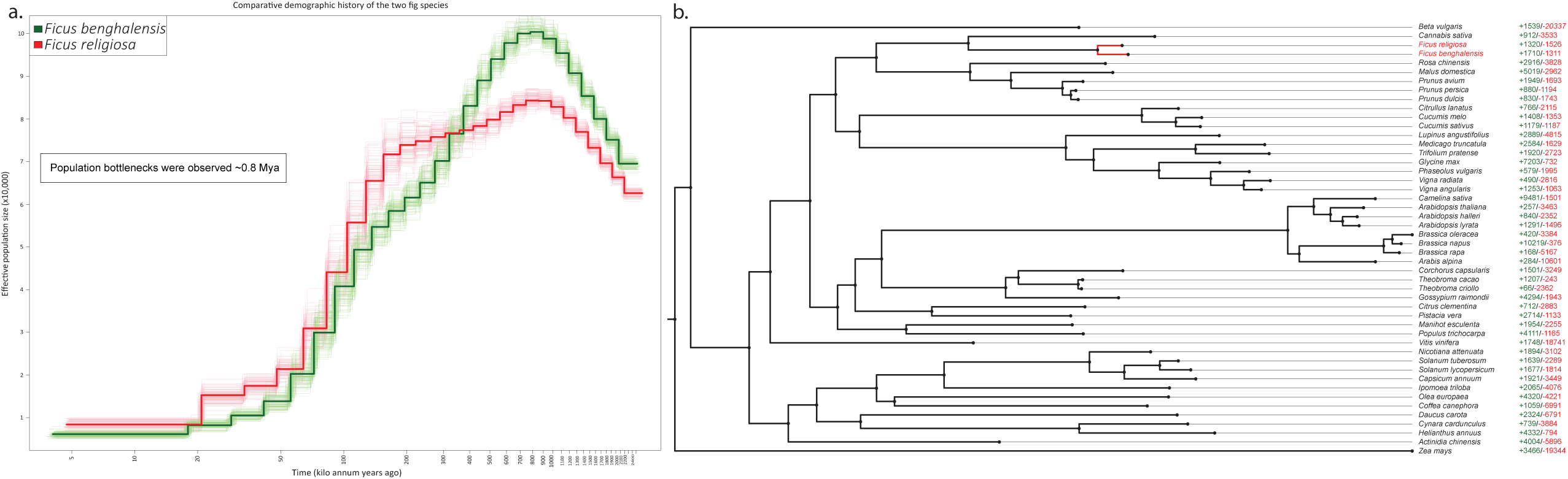

Phylogenetic position of *F. benghalensis* was resolved in comparison with 46 other Eudicot species spanning a total of 15 plant orders, along with *Zea mays* as an out-group species. A total of 507 fuzzy one-to-one orthogroups were constructed across all 48 selected species including *F. benghalensis* and *F. religiosa*, and maximum likelihood-based phylogenetic tree was constructed **(Figure 1b)** using the concatenated, filtered (for undetermined values) protein sequence alignments of these orthogroups, containing a total of 559,964 alignment positions.

*F. religiosa* and *F. benghalensis* that belong to the same genus and same plant family (Moraceae) were found to be the closest species of each other and were placed in the same clade. Four other species – *Rosa chinensis, Malus domestica, Prunus avium*, and *Cannabis sativa* from the same phylogenetic order Rosales were placed in the adjacent clades to the two *Ficus* species. Among these four species, *Cannabis sativa* belonging to the plant family Cannabaceae was the closest of *Ficus* species and shared the most recent common ancestor. *Ficus* plants belonging to Rosales plant order were closer to species from Fabales and Cucurbitales orders in comparison to other selected species, which is also supported by studies based on 18s rDNA, atpB and rbcL gene sequences^28^, and other previous reports^29^.

Relative phylogenetic positions of the other selected plant orders in this study were also supported by previous studies^28,30^. The phylogenetic positions of the plant orders – Rosales, Fabales and Cucurbitales were compared and found to be comparatively closer to the plant orders Brassicales, Malpighiales, Sapindales, and Malvales. The plant orders Apiales, Asterales, Solanales, Gentianales, Lamiales, Ericales, Vitales, and Caryophyllales diverged earlier, and among them species from the plant order Caryophyllales (*Beta vulgaris*) was found to diverge the earliest among all the selected Eudicot species, and hence it was the most distant Eudicot species to the two *Ficus* species.

### Expanded/contracted gene families

A total of 60,204 gene families were obtained using the coding gene sequences of the two *Ficus* species and 46 other plant species that were used for phylogenetic tree construction. 26,471 families were retained after the clade and size-based filtering of gene families. 1,710 families were expanded and 1,311 families were contracted in gene number in *F. benghalensis*. In *F. religiosa* species, 1,320 and 1,526 families were found to be expanded and contracted, respectively.

### Genes with signatures of adaptive evolution

A total of 7,468 orthogroups were identified across the 16 species including *F. benghalensis* and *F. religiosa* (see Methods) that were used for the evolutionary analyses to identify the *Ficus* species*-*specific genes showing signatures of adaptive evolution. Among the *F. benghalensis* species-specific genes, 1,449 genes had unique amino acid substitutions compared to the other plant species, and 1,001 of these genes were identified to have a functional impact. Also, 108 genes showed positive selection (with p-values < 0.05), and 26 genes showed higher rate of evolution i.e., higher branch length distance values compared to other species. Additionally, 268 genes containing positively selected codon sites were found with >95% probability, which is a site-specific evolutionary signature. A total of 45 genes showed at least two of the three evolutionary signatures – positive selection, higher rate of evolution, and unique substitution having functional impact, and thus were identified as the genes with Multiple Signs of Adaptive evolution (MSA). All these 45 genes were found to be associated with functions that are essential for long-time survival of plants e.g., root development, flowering and reproduction, cellular machinery, metabolism etc. **(Figure 2)**.

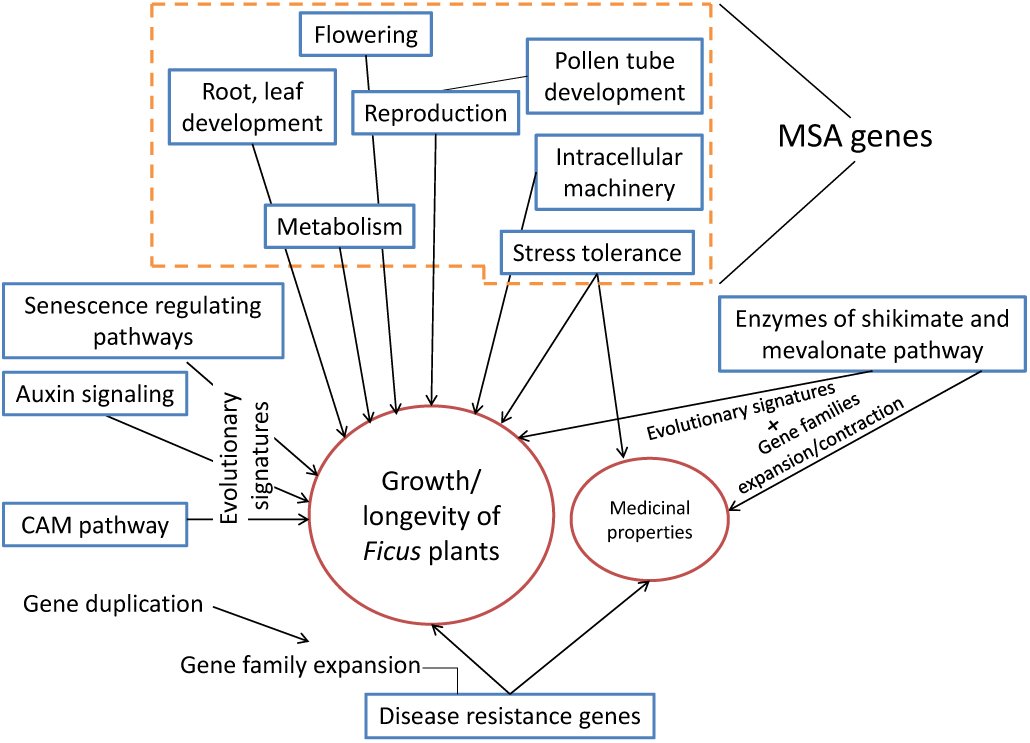

Similarly, in *F. religiosa* species a total of 111 genes were positively selected (with p-values < 0.05), and 30 genes showed higher rate of nucleotide divergence. Among the 1,360 genes that showed unique amino acid substitution, 978 genes were predicted to have functional impacts, and were considered for further analysis. Also, 255 genes possessed positively selected codon sites with >95% probability. A total of 55 genes were identified as MSA genes. All these 55 genes were found to be involved in functions that could be essential for providing longevity in plants such as root development, reproduction, seedling growth, metabolism etc. **(Figure 2)**.

### Enzymes involved in Shikimate and Mevalonate pathway

Shikimate and Mevalonate pathways are the two important pathways for fig pollination by production of VOCs, secondary metabolism, and plant development^7^. All seven enzymes involved in shikimate pathway and mevalonate pathway were present in the high-confidence coding gene set of *F. religiosa* and *F. benghalensis*. In *F. religiosa*, shikimate kinase, *DHQ* synthase (involved in shikimate pathway), and mevalonate kinase (involved in mevalonate pathway) showed unique amino acid substitution with functional impact compared to their orthologs in other 15 species. Furthermore, mevalonate pathway is also the precursor to terpenoid backbone biosynthesis. Two enzymes – *FPPS* (unique substitution with functional impact) and *GGPPS* (unique substitution with functional impact and positively selected codon sites) that are involved in the downstream terpenoid backbone biosynthesis pathway also showed evolutionary signatures. In *F. benghalensis, EPSPS* (involved in shikimate pathway) showed unique amino acid substitution with functional impact, and *GGPPS* (involved in terpenoid biosynthesis pathway) showed unique substitution with functional impact and positively selected codon sites.

Enzymes involved in mevalonate and shikimate pathway were also analyzed for gene family expansion/contraction. Gene families containing the genes of mevalonate pathway were not expanded/contracted in *F. benghalensis*, whereas one family was contracted (*MK*) in *F. religiosa*. For the gene families containing genes involved in shikimate pathway, three expanded (*DAHP* synthase, 3-dehydroquinate dehydratase, shikimate dehydrogenase) gene families were found in *F. benghalensis*, whereas one contracted family (shikimate kinase) was found in *F. religiosa*. Gene family expansion of the genes involved in shikimate pathway of *F. benghalensis* was accounted by gene duplication events. 3-dehydroquinate dehydratase and shikimate dehydrogenase genes were observed to be originated by tandem duplication, whereas DAHP synthase gene underwent segmental duplication.

## DISCUSSION

In this study, we have performed the first whole genome sequencing and analysis of *F. benghalensis* and *F. religiosa*, two dicot fig plants belonging to Moraceae family. *Ficus* plants are broadly known for their well-developed morphological characteristics as well as their ecological, evolutionary, medicinal, and religious importance. Thus, the whole genome sequences of these two *Ficus* species will serve as a valuable resource to study the significant characteristics of *Ficus* genus as well as Moraceae plant family. The hybrid sequencing and assembly approach using 10x Genomics linked reads and Oxford Nanopore long reads helped in successful construction of *F. benghalensis* and *F. religiosa* genomes with N50 values of 486.9 Kbp and 553.4 Kbp, respectively. A comparatively higher N50 value of *F. religiosa* genome assembly might be a result of lower heterozygosity (1.62%) and lower repeat content (51%) compared to *F. benghalensis* genome (1.78% heterozygosity and 54% repeat content). Higher BUSCO completeness and higher percentage of reads mapped to the genome assembly also attests to the genome assembly quality.

The genome size of these two *Ficus* species is similar to *F. microcarpa* genome and slightly larger than other *Ficus* - *F. carica, F. hispida* and *F. erecta* genomes^7,18,19^. However, the estimated and assembled genome sizes of the *Ficus* plants in this study is smaller than their genome sizes reported at Plant C-values Database, which was obtained using experimental methodologies that depend on various factors such as the usage of reference standard plant species while estimating the genome size^31^. Thus, in this study, three different approaches - GenomeScope **(Supplementary Figures 1a-1b)**, Supernova assembler, and SGA-preqc were used to estimate the genome sizes of both the *Ficus* species that provide additional confirmations on smaller genome sizes compared to the sizes reported at Plant C-values Database.

The final genome assemblies were used for coding gene prediction along with transcriptome and protein sequences as empirical evidence, and AED value criteria of <0.5 was used as quality control metric to construct the high-confidence gene set. Presence of 94.4% (complete and fragmented) and 94.3% (complete and fragmented) BUSCO genes and all the genes present in shikimate and mevalonate pathway in this high-confidence gene set attests to the comprehensiveness of the coding gene set of *F. benghalensis* and *F. religiosa*, respectively. The number of high-confidence coding genes is also similar to *F. hispida* and *F. microcarpa* genomes^7^. Further, the observed high percentage of duplicated genes in these genomes may have occurred due to expansion of gene families that provide long lifespan to plants, as in case of Oak genome^32^. Also, the high percentage of inter-species collinearity between the two *Ficus* genomes might be a result of *Ficus* hybridization events caused by frequent host switching of the fig pollinators^33^. Among the LTR retrotransposon class of repeats, Gypsy/DIRS1 elements were higher than Ty1/Copia elements in both the species (**Supplementary Tables 5-6**), similar to other *Ficus* genomes^7,18,34^.

The comprehensive genome-wide phylogeny including 48 species (with two fig plants) covering 15 phylogenetic plant orders constructed using a large number of one-to-one orthogroups and a decent bootstrap value (no polytomy was observed) will serve as a valuable reference for future phylogenetic studies. The predicted population bottleneck events in case of both these *Ficus* plants might be the result of major glaciation events in Pleistocene^35^, and the environmental changes (warm and cold conditions prevailing for longer time, lack of rainfall etc.) caused in the Middle Pleistocene transition (MPT)^36,37^. PSMC analysis also showed that these *Ficus* species could not recover from the long-term decrease in effective population size caused by the bottleneck events.

The inclusion of closely related plant species (according to the phylogenetic tree) for comparative analysis helped in precise identification of genes with evolutionary signatures in *F. benghalensis* and *F. religiosa* species. Further the inclusion of coding genes from both the fig plants in this analysis helped to identify plausible species-specific *Ficus* genes responsible for the morphological characteristics and evolutionary adaptation in these two plant species.

Evolutionary signatures as well as gene family expansion/contraction for the enzymes involved in shikimate pathway in *F. benghalensis*, and for the enzymes involved in both shikimate and mevalonate pathways in case of *F. religiosa* suggest adaptive evolution of these pathways. Evolution of shikimate pathway, which is also responsible for the production of pollinator attracting VOCs^7^, suggest presence of fig-wasp obligate mutualism in these plant species. Similarly, the evolution of mevalonate pathway in *F. religiosa* that regulates plant developmental processes by aiding metabolism, phytohormone biosynthesis, isoprenoid biosynthesis etc.^38,39^, indicates the adaptive evolution of plant growth and development in this species. In addition, the adaptive evolution of shikimate and mevalonate pathways, which are also the precursors for production of phenolic and terpenoid classes of secondary metabolites^40,41^, explain the presence of medicinal properties in these plants. To support this, evolutionary signatures were also observed in other enzymes involved in terpenoid backbone biosynthesis pathway in both the species.

Genes that are associated with providing longevity were identified with signatures of adaptive evolution in both the species. Among the MSA genes related to growth and development, the genes involved in plant root development, flowering, vegetative growth, and metabolism were notable. Flowering related MSA genes in both the species also might influence fig pollination as well as fig-wasp mutualism and co-evolution. The study also reports the presence of CAM pathway of photosynthesis in *F. benghalensis* and *F. religiosa*, which is a characteristic of epiphyte or hemi epiphyte plants to conserve water storage, when they survive as epiphytes^42^. Genes involved in CAM pathway showed signatures of adaptive evolution in both these species. Studies have shown that plants that use both C_3_ and CAM pathway of photosynthesis exhibit long-lived leaves, thus the evolutionary signatures of CAM pathway in *F. benghalensis* and *F. religiosa* perhaps also contribute to leaf longevity^43^.

84% and 91% of the MSA genes were also associated with tolerance against biotic and abiotic stress responses in *F. benghalensis* and *F. religiosa*, respectively, which further assist in plant survival against environmental challenges. Presence of diverse defense or stress tolerance-related genes is also an important factor in providing long-time survival of plants^44^. Stress response mechanisms such as JA and SA-signaling pathways also regulate plant secondary metabolism^45^, which is further related to the medicinal properties of plants^46^. Taken together, it can be speculated that genes that help to survive in diverse and challenging environmental conditions e.g., genes involved in plant growth and development (root cell growth and proliferation, auxin signaling, CAM pathway, plant senescence regulating pathways, plant metabolism, cell wall and plasma membrane component biosynthesis, intracellular machinery) have evolved to provide longer life-span in plant species like *F. benghalensis* and *F. religiosa*, which are keystone plant species in tropical and sub-tropical ecosystems^2,5^.

## METHODS

### DNA-RNA extraction and Sequencing

The leaves were collected from a *F. benghalensis* tree and a *F. religiosa* tree located in Bhopal, India (23.2280252°N 77.2088987°E). The leaves were cleaned and processed for DNA and RNA extraction. The DNA extraction was carried out using Carlson lysis buffer by following Oxford Nanopore suggested protocol using Genomic tip 20 of Blood and cell culture kit (Qiagen, CA, USA) with few modifications. The quality check and quantification of DNA was performed on Nanodrop 8000 spectrophotometry (Thermo Scientific) and Qubit 2.0 flourometer (Invitrogen, Thermofisher Scientific, USA). The species was identified as *F. benghalensis* and *F. religiosa* by amplifying, sequencing and performing Nucleotide BLAST (Basic local Alignment Search Tool) of two marker genes i.e., Internal Transcribed Spacer (ITS) and Maturase K (MatK). The genomic library for linked reads was prepared on Chromium Controller instrument (10X Genomics, CA) and sequenced on Illumina Novaseq 6000 platform to produce 150 bp paired-end reads. The long read genomic library was prepared using SQK-LSK-109 kit followed by sequencing on MinION instrument (Oxford Nanopore Technologies, UK). The total RNA was extracted using TriZol reagent (Invitrogen, Thermofisher Scientific, USA). The transcriptomic library was prepared using Illumina TruSeq Stranded Total RNA library preparation kit with Ribo-Zero workflow and 150 bp paired-end reads were generated on Illumina Novaseq 6000 instrument (Illumina Inc., CA, USA). The detailed information related to species identification and DNA-RNA extraction is mentioned in **Supplementary Notes 1**.

### Genome assembly

Genome sizes of the two Ficus plant species were predicted using 10x Genomics linked reads. Barcode sequences were extracted from the raw reads using available proc10xG set of python scripts (https://github.com/ucdavis-bioinformatics/proc10xG), and these pre-processed linked reads were used to estimate the genome size using a k-mer distribution-based approach executed by SGA-preqc^20^(in paired-end mode). Genomic heterozygosity was also estimated for the two Ficus species in order to assess the complexity of the genomes. Jellyfish^47^v2.2.10 was used to generate k-mer count histograms using the barcode-filtered paired-end reads, and GenomeScope^22^v2.0 was used to calculate the heterozygosity profiles.

The de novo genome assembly of both the Ficus species was performed following the same approach. Oxford Nanopore raw data was basecalled using Guppy v3.2.1 (Oxford Nanopore Technologies) and adapter removal of the raw reads were performed using Porechop v0.2.3 (Oxford Nanopore Technologies). Genome assembly using these pre-processed reads was performed using wtdbg v2.0.0^48^with default settings, SMARTdenovo (https://github.com/ruanjue/smartdenovo) with minimum read length of 0, Canu v2.0^49^with default settings (after reads correction using Canu v2.0) and Flye v2.4.2^50^with default settings. Quality statistics of all these assemblies were calculated using Quast v5.0.2^51^, and based on the assembled genome size and assembly statistics, the assembly derived from Flye v2.4.2 was selected for further analysis. 10x Genomics linked reads, which were processed using Longranger basic v2.2.0 (https://support.10xgenomics.com/genome-exome/software/pipelines/latest/installation), were used to polish this assembly three times using Pilon v1.23^52^. RNA-Seq data was quality-filtered using Trimmomatic^53^v0.38, and the filtered paired-end data was used for scaffolding using AGOUTI v0.3.3^54^, to improve the assembly contiguity. The barcode-processed linked reads and pre-processed Nanopore long reads (Canu v2.0-corrected reads) were also used for further scaffolding of this assembly using ARCS v1.1.1^55^and LINKS v1.8.6^56^with default settings. Contigs with length of ≥1,000 scaffolding) were extracted (assembly v0.1).

A total of 327.41 million (116.67x raw reads coverage) and 161.11 million (~56x raw reads coverage) barcoded 10x Genomics linked reads were used for de novo assembly of *F. benghalensis* genome using Supernova v2.1.1^21^. For *F. religiosa* genome assembly, a total of 324.53 million reads (125.43X raw coverage) and 154.24 million reads (59X raw coverage) were used in Supernova v2.1.1. The assemblies generated using 327.41 million barcoded linked reads (116.67x raw coverage) for F. benghalensis, and 324.53 million 10x Genomics linked reads (125.43X raw coverage) for *F. religiosa* were considered for downstream analysis based on better assembly quality parameters. The mis-assembly regions were corrected using Tigmint v1.2.1^57^with the barcode-processed linked reads resulted from Longranger basic v2.2.0. Quality-filtered RNA-Seq paired-end reads were used to improve the contiguity of this assembly using AGOUTI v0.3.3^54^. Barcode-processed linked reads, and error-corrected Nanopore long reads were used for further scaffolding using ARCS v1.1.1^55^(default settings) and LINKS v1.8.6^56^, respectively. Contigs with length of ≥1,000 bp (after scaffolding) were retained (assembly v0.2).

For each of the two species, assembly v0.1 and assembly v0.2 were merged using Quickmerge v0.3^58^, which joins the gaps between the sequences in one assembly by using sequences from the other assembly, and thus improve the assembly contiguity (assembly v1). Further, contigs from assembly v0.2 were searched against the contigs from assembly v1 using BLASTN, and the unique sequences that did not match even with stringent parameters **(Supplementary Notes 2)** were directly added to assembly v1^59,60^. Gap-closing of these assemblies (assembly v2) was performed in three iterations using LR_Gapcloser^61^ with error-corrected Oxford Nanopore long reads. Finally, barcode-processed 10x Genomics linked reads were used to fix any further local mis-assemblies, small indels, and base errors using Pilon v1.23^52^, to generate the final *F. benghalensis* and *F. religiosa* genome assemblies (assembly v3). Completeness of these genome assemblies was checked using embryophyta_odb10 single-copy orthologs dataset from BUSCO v5.2.1^62^. Also, to check the comprehensiveness of the final genome assembly, barcode-filtered 10x Genomics linked reads and adapter-processed Nanopore long reads were mapped to the final genome assemblies using BWA-MEM^63^ v0.7.17, and the mapping percentage was calculated using SAMtools^64^v1.9 “flagstat”. The detailed information related to genome assembly and processing steps are mentioned in **Supplementary Figure 2 and Supplementary Notes 2**.

Sequence variation in the final Ficus genome assemblies was analyzed by mapping the barcode-filtered 10x Genomics linked reads to their respective whole genome assemblies (assembly v3) using BWA-MEM^63^ v0.7.17, and SAMtools^64^ v1.9. Variant calling was performed using BCFtools^65^ v1.9 “mpileup” with the quality-filtration criteria - sequencing depth ≥30, quality of variant sites ≥30, and mapping quality ≥50.

### Genome annotation and Gene set construction

Genome annotation for *F. benghalensis* and *F. religiosa* species was performed using their corresponding final genome assemblies (assembly v3) generated in the previous step. For repeat annotation, a de novo repeat library was constructed for these genomes using RepeatModeler v2.0.1^66^, which was further clustered to remove redundant sequences using CD-HIT-EST v4.8.1^67^ with sequence identity of 90%, and seed size of 8 bp. This resultant repeat library was used to soft-mask the two Ficus genomes using RepeatMasker v4.1.0 (http://www.repeatmasker.org). Additionally, simple repeat sequences were also predicted in the final genome assemblies (assembly v3) using Tandem Repeats Finder (TRF) v4.09^23^ **(Supplementary Notes 2)**.

The repeat-masked genomes of *F. benghalensis* and *F. religiosa* were used for gene set construction. MAKER v2.31.10^24^, the genome annotation pipeline was used to construct the coding gene sets using ab initio and evidence alignment-based approaches. Quality-filtered RNA-Seq reads (using Trimmomatic v0.38) of *F. benghalensis* from this study, and quality-filtered transcriptome reads of *F. religiosa* from this study and previous report^68^ were used to construct de novo transcriptome assemblies using Trinity v2.9.1^69^ with default parameters **(Supplementary Notes 2)**. The resultant transcriptome assemblies, and the protein sequences of phylogenetically closer species - Cannabis sativa, Malus domestica, Prunus avium, and Rosa chinensis that belong to the same plant order Rosales (obtained from Ensembl Plants release 48) were used as EST and protein evidence for gene set construction of the two respective Ficus plant species. For ab initio gene finding, AUGUSTUS v3.2.3^70^ and were used, and for empirical evidence alignments, BLAST^71^ and EXONERATE v2.2.0 (https://github.com/nathanweeks/exonerate) were used. The gene models were filtered based on their Annotation Edit Distance (AED) values and length-based criteria. Only the genes with AED values <0.5, and length (≥300 bp) were selected for the final gene set construction of *F. benghalensis* and *F. religiosa*, and termed as high-confidence gene sets^72^. Completeness of these gene sets was assessed using BUSCO v5.2.1^62^ with embryophyta_odb10 dataset of single-copy orthologous genes.

Further, de novo prediction of tRNA and rRNA genes was performed using tRNAscan-SE v2.0.7^73^, and Barrnap v0.9 (https://github.com/tseemann/barrnap). Homology-based identification of miRNA gene sequences was performed using miRBase database^74^ **(Supplementary Notes 2)**.

### PSMC analysis

Demographic history was constructed for *F. benghalensis* and *F. religiosa* plant species for comparative study using Pairwise Sequentially Markovian Coalescent (PSMC)^26^ with the whole genome sequences of the corresponding Ficus species. The barcode-filtered 10x Genomics linked reads of the two species, which were mapped to their respective whole genome assemblies (assembly v3) using BWA-MEM^63^, were used to construct the consensus diploid sequences using SAMtools v1.9^64^ “mpileup”, and BCFtools v1.9^65^ in a manner that the minimum read depth is a third of the average read depth, and the maximum read depth is set to the twice of the average read depth. The consensus sequences were filtered for sites with quality score <20. PSMC analysis was performed using the parameters “N30 -t5 -r5 -p4+25*2+4+6” with 100 bootstrap values. Generation time of 7 years, and a mutation rate per site per generation of 1.75e-08 was considered for this analysis, which was used for another Ficus species F. carica in a previous study^27^.

### Phylogenetic analysis

Protein sequences of all 45 Eudicot species, available on Ensembl^75^ plants release 48, were used to resolve the phylogenetic position of *F. benghalensis* and *F. religiosa*. The selected Eudicot species were – *Actinidia chinensis, Arabidopsis halleri, Arabidopsis lyrata, Arabidopsis thaliana, Arabis alpina, Beta vulgaris, Brassica napus, Brassica oleracea, Brassica rapa, Camelina sativa, Cannabis sativa female, Capsicum annuum, Citrullus lanatus, Citrus clementina, Coffea canephora, Corchorus capsularis, Cucumis melo, Cucumis sativus, Cynara cardunculus, Daucus carota, Glycine max, Gossypium raimondii, Helianthus annuus, Ipomoea triloba, Lupinus angustifolius, Malus domestica Golden, Manihot esculenta, Medicago truncatula, Nicotiana attenuata, Olea europaea var. sylvestris, Phaseolus vulgaris, Pistacia vera, Populus trichocarpa, Prunus avium, Prunus persica, Prunus dulcis, Rosa chinensis, Solanum lycopersicum, Solanum tuberosum, Theobroma cacao Belizian Criollo B97-61/B2, Theobroma cacao Matina 1-6, Trifolium pratense, Vigna angularis, Vigna radiata, and Vitis vinifera*. Proteome files of these 45 Eudicot species along with MAKER-derived protein sequences of *F. benghalensis* and *F. religiosa* were used for phylogenetic analysis. *Zea mays*, which is a monocot plant, was used as an outgroup species, and protein sequences of *Zea mays* were also obtained from Ensembl plants release 48.

Proteome files of all species were used to extract the longest isoform for each protein. The resultant protein sequences were used to construct the orthologous gene sets using OrthoFinder v2.4.1^76^. From these orthogroups, the fuzzy one-to-one orthogroups that contained proteins from all 48 species were extracted using KinFin v1.0^77^. For further analysis, these fuzzy one-to-one orthologous gene sets were processed to extract and consider only the longest protein sequence for each species. Each of these orthogroups was individually aligned using MAFFT v7.467^78^. These multiple sequence alignments were filtered to remove the empty sites and were concatenated using BeforePhylo v0.9.0 (https://github.com/qiyunzhu/BeforePhylo). RAxML v8.2.12^79^, which uses rapid hill climbing algorithm, was utilized to construct the maximum likelihood-based species phylogenetic tree with the concatenated protein sequence alignment. ‘PROTGAMMAAUTO’ amino acid substitution model and a bootstrap value of 100 were used for phylogenetic tree construction.

### Evolution of gene families

CAFÉ v4.2.1^80^ was used to estimate the evolution of gene families across all 48 selected species (Zea mays as an outgroup species) using the proteome files containing the longest isoform for each protein, and the species phylogenetic tree generated in the previous step. Prior to the analysis, the species phylogenetic tree (with *Zea mays* as an outgroup species) was converted to ultrametric tree based on the calibration point of 121 Million years between *F. benghalensis*, and *Beta vulgaris*, obtained from TimeTree database^81^. All-versus-All BLASTP search for the protein sequences of all 48 species was performed, which was followed by clustering using MCL v14.137^82^, and removal of gene families that contained genes in <2 species of the specified clades and ≥100 gene copies for at least one species. These filtered gene families, and the ultrametric species tree was utilized to analyze the expansion or contraction of gene families using two-lambda (λ) model, where λ is a random birth-death parameter. In this analysis, *F. benghalensis, F. religiosa*, and their phylogenetically closer species – *Cannabis sativa, Rosa chinensis, Malus domestica, Prunus avium, Prunus persica*, and *Prunus dulcis* were given the same λ-value, while the other species were given another λ-value. The considerably expanded (with ≥10 expanded genes) and contracted (with ≥5 contracted genes) gene families for *F. benghalensis* and *F. religiosa* were extracted and compared.

Further, gene family expansion/contraction for the enzymes involved in mevalonate and shikimate pathway that play important role in *Ficus* plants^7^ were analyzed using CAFÉ results. EC numbers of these enzymes were used to extract their sequence (specific to *Arabidopsis thaliana*, a model Eudicot plant) from Swiss-Prot database^83^, and the sequences were queried against the proteome of *F. benghalensis* and *F. religiosa* using BLASTP (e-value of 10^−5^) to identify the enzymes in these two *Ficus* species. Gene families of these enzymes were identified from CAFÉ results, and the expansion/contraction of these families was identified in two *Ficus* species compared to their immediate ancestor phylogenetic node.

### Synteny analysis

Collinear blocks between *F. benghalensis* and *F. religiosa* species (inter-species), and intra-species syntenic blocks for these two species were identified using the All-versus-All BLASTP (e-value of 10^−5^) homology alignments of the respective protein sequences and the GFF annotations. Inter-species and intra-species synteny analysis was performed using MCScanX^25^, 10 genes were required to detect a collinear block^84^. Also, the duplicated genes in each of the Ficus species were detected and classified into various gene duplication categories (proximal, tandem, segmental, or dispersed) using the same parameter.

### Identification of Evolutionary signatures

For identifying the genes with evolutionary signatures, only the Eudicot species from the phylogenetically closer plant orders of the two *Ficus* species (one species from each order) were considered to avoid erroneous results and to analyze adaptive evolution of these two *Ficus* species across dicot plant orders. The selected 14 Eudicot species were - *Arabidopsis thaliana* (order Brassicales), *Actinidia chinensis* (order Ericales), *Coffea canephora* (order Gentianales*), Cucumis melo* (order Cucurbitales), *Daucus carota* (order Apiales), *Glycine max* (order Fabales), *Gossypium raimondii* (order Malvales), *Helianthus annuus* (order Asterales), *Olea europaea var. sylvestris* (order Lamiales), *Pistacia vera* (order Sapindales), Populus trichocarpa (order Malpighiales), *Rosa chinensis (*order Rosales), *Solanum tuberosum* (order Solanales), and *Vitis vinifera* (order Vitales). Proteome files of these 14 species along with that of *F. benghalensis* and *F. religiosa* were used to construct the orthologous gene sets across 16 species using OrthoFinder v2.4.1^76^. Orthogroups containing protein sequences from all 16 species were identified, and only the longest protein sequence for each species was extracted from these orthogroups.

### Genes with unique amino acids substitution

Each of the orthogroups across 16 species constructed in the previous step was aligned using MAFFT v7.467^78^. In the multiple sequence alignments, *F. benghalensis* genes showing amino acid positions that were identical in other species but different in this species were considered as genes with unique amino acid substitutions in *F. benghalensis*. Similarly, *F. religiosa* genes that showed different amino acids in the sequence alignments compared to the other species were also considered as the *F. religiosa* genes with unique substitutions. Any gap in these alignments and ten sites around the gap were not considered in this analysis. The genes showing unique amino acid substitutions were extracted, and impact of these unique amino acid substitutions on the proteins function was predicted using Sorting Intolerant From Tolerant (SIFT)^85^ with UniProt reference database^86^.

### Genes with higher nucleotide divergence

The protein sequence alignments of the orthogroups across 16 species generated in the previous step were used to construct individual maximum likelihood-based phylogenetic tree for each orthogroup using RAxML v8.2.12^79^ with ‘PROTGAMMAAUTO’ amino acid substitution model and 100 bootstrap values. Root-to-tip branch length distances were estimated for genes from all selected species in these phylogenetic trees using “adephylo”^87^ package in *R. F. benghalensis* and *F. religiosa* specific genes with relatively higher branch length distance values compared to the other species were extracted separately as the genes with higher rate of evolution.

### Positively selected genes

The nucleotide sequences of all the orthologous gene sets constructed across 16 species were individually aligned using MAFFT v7.467^78^. Further, RAxML^79^ v8.2.12 was used to construct the species phylogenetic tree for these selected 16 species in a similar way as mentioned in the earlier steps (see Phylogenetic analysis step). This species phylogenetic tree and the nucleotide alignments for the orthogroups in phylip format were used to separately detect the *F. benghalensis* and *F. religiosa* genes with positive selection using a branch-site model utilized by “codeml” program available in PAML v4.9a^88^. Likelihood-ratio tests were performed from the obtained log-likelihood values and genes qualifying against the null model having FDR-corrected p-values of <0.05 (obtained from chi-square analysis) were termed as positively selected genes. Additionally, Bayes Empirical Bayes (BEB) analysis was carried out to identify the codon sites of *F. benghalensis* and *F. religiosa* genes that showed positive selection with >95% probability for the foreground lineage.

The genes that showed more than one of these above mentioned evolutionary signatures, namely - positive selection, higher nucleotide divergence rate, and unique substitution with functional impact, were extracted and considered as the genes with multiple signatures of adaptive evolution (MSA genes)^89,90^.

### Functional annotation

The complete high-confidence gene sets of *F. benghalensis* and *F. religiosa* was annotated against NCBI Non-Redundant (nr) database using BLASTP (e-value of 10^−5^), Swiss-Prot^83^ database using BLASTP (e-value of 10^−5^), and Pfam-A v32.0^91^ database using HMMER v3.3^92^ (e-value of 10^−5^). The species-specific gene sets with evolutionary signatures were assigned to KEGG Orthology (KO) categories, and KEGG pathways using KAAS^93^ v2.1 genome annotation server. These genes were also annotated to different COG categories using eggNOG-mapper v2^94^. Gene Ontology (GO) categories were also assigned to these genes using WebGeStalt^95^ web server, and in the over-representation analysis, only the GO categories with p-values <0.05 were retained. Further, functional annotations of these gene sets were screened, and curated manually. The MSA genes were also checked for protein-protein interaction data in STRING v11.0 database^96^ using information from previously reported studies, and the interacting genes were analyzed. Enzymes involved in shikimate pathway and Eukaryotic mevalonate pathway were searched for their presence in the gene sets showing evolutionary signatures, and in the expanded/contracted gene families.

## ACKNOWLEDGEMENTS

AC, SM, and MSB thank Council of Scientific and Industrial Research (CSIR) for fellowship. The authors also thank the NGS facility at IISER Bhopal and the intramural research funds provided by IISER Bhopal.

## AUTHORS’ CONTRIBUTIONS

VKS conceived and coordinated the project. SM performed DNA-RNA extraction, prepared the samples for sequencing, performed the Nanopore sequencing, and the species identification assays. AC and VKS designed the computational framework of the study. AC performed all the computational analysis presented in the study, performed the functional annotation of gene sets, and constructed the figures. AC, MSB and VKS analyzed the data and interpreted the results. AC, SM and VKS wrote the manuscript. All the authors have read and approved the final version of the manuscript.

