## Supplementary Information for "Genome sequencing and comparative analysis of *Ficus benghalensis* and *Ficus religiosa* trees reveal evolutionary mechanisms of longevity"

### SUPPLEMENTARY TABLES

**Supplementary Table 1. Summary of the genomic data generated for *Ficus* genomes**

| Species | Total 10x Genomics reads | Total 10x Genomics data (bases) | Total Nanopore reads | Total Nanopore data (bases) |
|---|---|---|---|---|
| *Ficus benghalensis* | 163,707,315 (x2) | 49,015,662,174 | 3,416,392 | 9,015,677,442 |
| *Ficus religiosa* | 162,262,671 (x2) | 48,681,137,914 | 2,612,589 | 7,186,698,379 |

Note: The sequencing coverage was calculated based on the genome sizes estimated using SGA-preqc[1]

**Supplementary Table 2. Summary of the transcriptome data generated for *Ficus* genomes**

| Species | Average read length R1 (bases) | Average read length R2 (bases) | Total Number of Read pairs | Total Number of Bases |
|---|---|---|---|---|
| *F. benghalensis* | 139.52 | 139.55 | 26,570,361 (x2) | 7,415,158,369 |
| *F. religiosa* | 135.47 | 135.49 | 39,690,127 (x2) | 10,754,429,937 |

**Supplementary Table 3. Summary statistics of the *de novo* genome assemblies**

| Parameters | *F. benghalensis* genome assembly | *F. religiosa* genome assembly |
|---|---|---|
| Number of contigs | 4,822 | 6,087 |
| Number of contigs (>= 5,000 bp) | 3,075 | 3,402 |
| Number of contigs (>= 10,000 bp) | 2,407 | 2,505 |
| Number of contigs (>= 25,000 bp) | 1,613 | 1,464 |
| Number of contigs (>= 50,000 bp) | 1,090 | 854 |
| Total length (bp) | 392,886,432 | 332,972,653 |
| Total length (>= 5,000 bp) | 388,855,687 | 326,957,949 |
| Total length (>= 10,000 bp) | 384,009,904 | 320,580,625 |
| Total length (>= 25,000 bp) | 371,230,410 | 303,781,488 |
| Total length (>= 50,000 bp) | 352,658,314 | 282,216,944 |
| Largest contig (bp) | 5,391,251 | 4,514,195 |
| N50 value (bp) | 486,949 | 553,438 |
| GC % | 34.54 | 34.31 |
| L50 value | 185 | 146 |
| Number of N's per 100 kbp | 230.90 | 918.02 |

**Supplementary Table 4. BUSCO statistics of the *de novo* genome assemblies**

| Parameters | *F. benghalensis* genome assembly | *F. religiosa* genome assembly |
|---|---|---|
| Complete BUSCOs (C) | 1,555 (96.4%) | 1,543 (95.6%) |
| Fragmented BUSCOs (F) | 27 (1.7%) | 17 (1.1%) |
| Missing BUSCOs (M) | 32 (1.9%) | 54 (3.3%) |
| Total BUSCO groups searched | 1,614 | 1,614 |

**Supplementary Table 5. Summary statistics of the *F. benghalensis* repetitive genomics regions detected by RepeatMasker**

| | | | | | |
|---|---|---|---|---|---|
| Total length: | 392,886,432 bp | | | | |
| GC (%) | 34.54% | | | | |
| Bases masked: | 201,773,676 bp (51.36%) | | | | |
| | | | Number of Elements | Length occupied | Percentage of Sequence |
| Retroelements | | | 47,893 | 46,407,496 bp | 11.81% |
| | SINEs | | 241 | 41,209 bp | 0.01% |
| | Penelope | | 0 | 0 bp | 0.00% |
| | LINEs | | 7,152 | 2,634,863 bp | 0.67% |
| | | CRE/SLACS | 0 | 0 bp | 0.00% |
| | | L2/CR1/Rex | 561 | 268,428 bp | 0.07% |
| | | R1/LOA/Jockey | 103 | 19,617 bp | 0.00% |
| | | R2/R4/NeSL | 0 | 0 bp | 0.00% |
| | | RTE/Bov-B | 0 | 0 bp | 0.00% |
| | | L1/CIN4 | 6,488 | 2,346,818 bp | 0.60% |
| | LTR elements: | | 40,500 | 43,731,424 bp | 11.13% |
| | | BEL/Pao | 1,205 | 170,004 bp | 0.04% |
| | | Ty1/Copia | 11,552 | 10,626,612 bp | 2.70% |
| | | Gypsy/DIRS1 | 27,509 | 32,540,804 bp | 8.28% |
| | | Retroviral | 0 | 0 bp | 0.00% |
| DNA transposons | | | 16,145 | 9,376,659 bp | 2.39% |

| | | | | | |
|---|---|---|---|---|---|
| | hobo-Activator | | 2,324 | 1,014,808 bp | 0.26% |
| | Tc1-IS630-Pogo | | 0 | 0 bp | 0.00% |
| | En-Spm | | 0 | 0 bp | 0.00% |
| | MuDR-IS905 | | 0 | 0 bp | 0.00% |
| | PiggyBac | | 0 | 0 bp | 0.00% |
| | Tourist/Harbinger | | 5,630 | 1,800,586 bp | 0.46% |
| | Other (Mirage, P-element, Transib) | | 0 | 0 bp | 0.00% |
| Rolling-circles | | | 13,215 | 4,169,792 bp | 1.06% |
| Unclassified: | | | 461,435 | 129,187,311 bp | 32.88% |
| Total interspersed repeats: | | | | 184,971,466 bp | 47.08% |
| Small RNA: | | | 8,950 | 1,538,441 bp | 0.39% |
| Satellites: | | | 0 | 0 bp | 0.00% |
| Simple repeats: | | | 253,235 | 9,047,865 bp | 2.30% |
| Low complexity: | | | 40,261 | 2,046,112 bp | 0.52% |

**Supplementary Table 6. Summary statistics of the *F. religiosa* repetitive genomics regions detected by RepeatMasker**

| | | | | | |
|---|---|---|---|---|---|
| Total length: | 332,972,653 bp | | | | |
| GC (%) | 34.31% | | | | |
| Bases masked: | 152,770,947 (45.88%) | | | | |
| | | | Number of Elements | Length occupied | Percentage of Sequence |

| | | | | | |
|---|---|---|---|---|---|
| Retroelements | | | 28,495 | 22,301,689 bp | 6.70% |
| | SINEs | | 1,053 | 285,093 bp | 0.09% |
| | Penelope | | 0 | 0 bp | 0.00% |
| | LINEs | | 4,714 | 1,428,760 bp | 0.43% |
| | | CRE/SLACS | 0 | 0 bp | 0.00% |
| | | L2/CR1/Rex | 0 | 0 bp | 0.00% |
| | | R1/LOA/Jockey | 0 | 0 bp | 0.00 % |
| | | R2/R4/NeSL | 0 | 0 bp | 0.00% |
| | | RTE/Bov-B | 0 | 0 bp | 0.00 % |
| | | L1/CIN4 | 4,714 | 1,428,760 bp | 0.43% |
| | LTR elements: | | 22,728 | 20,587,836 bp | 6.18% |
| | | BEL/Pao | 0 | 0 bp | 0.00% |
| | | Ty1/Copia | 7,375 | 5,176,206 bp | 1.55% |
| | | Gypsy/DIRS1 | 14,955 | 15,015,388 bp | 4.51% |
| | | Retroviral | 0 | 0 bp | 0.00 % |
| DNA transposons | | | 10,104 | 5,446,639 bp | 1.64% |
| | hobo-Activator | | 3,623 | 719,343 bp | 0.22% |
| | Tc1-IS630-Pogo | | 117 | 30,151 bp | 0.01% |
| | En-Spm | | 0 | 0 bp | 0.00% |
| | MuDR-IS905 | | 0 | 0 bp | 0.00% |
| | PiggyBac | | 0 | 0 bp | 0.00% |
| | Tourist/Harbinger | | 913 | 489,268 bp | 0.15% |

| | | | | | |
|---|---|---|---|---|---|
| | Other (Mirage, P-element, Transib) | | 0 | 0 bp | 0.00% |
| Rolling-circles | | | 4,497 | 2,079,581 bp | 0.62% |
| Unclassified: | | | 406,515 | 111,031,431 bp | 33.35% |
| Total interspersed repeats: | | | | 138,779,759 bp | 41.68% |
| Small RNA: | | | 2,611 | 979,394 bp | 0.29% |
| Satellites: | | | 965 | 175,602 bp | 0.05% |
| Simple repeats: | | | 252,720 | 8,792,487 bp | 2.64% |
| Low complexity: | | | 39,919 | 1,964,124 bp | 0.59% |

### SUPPLEMENTARY FIGURES

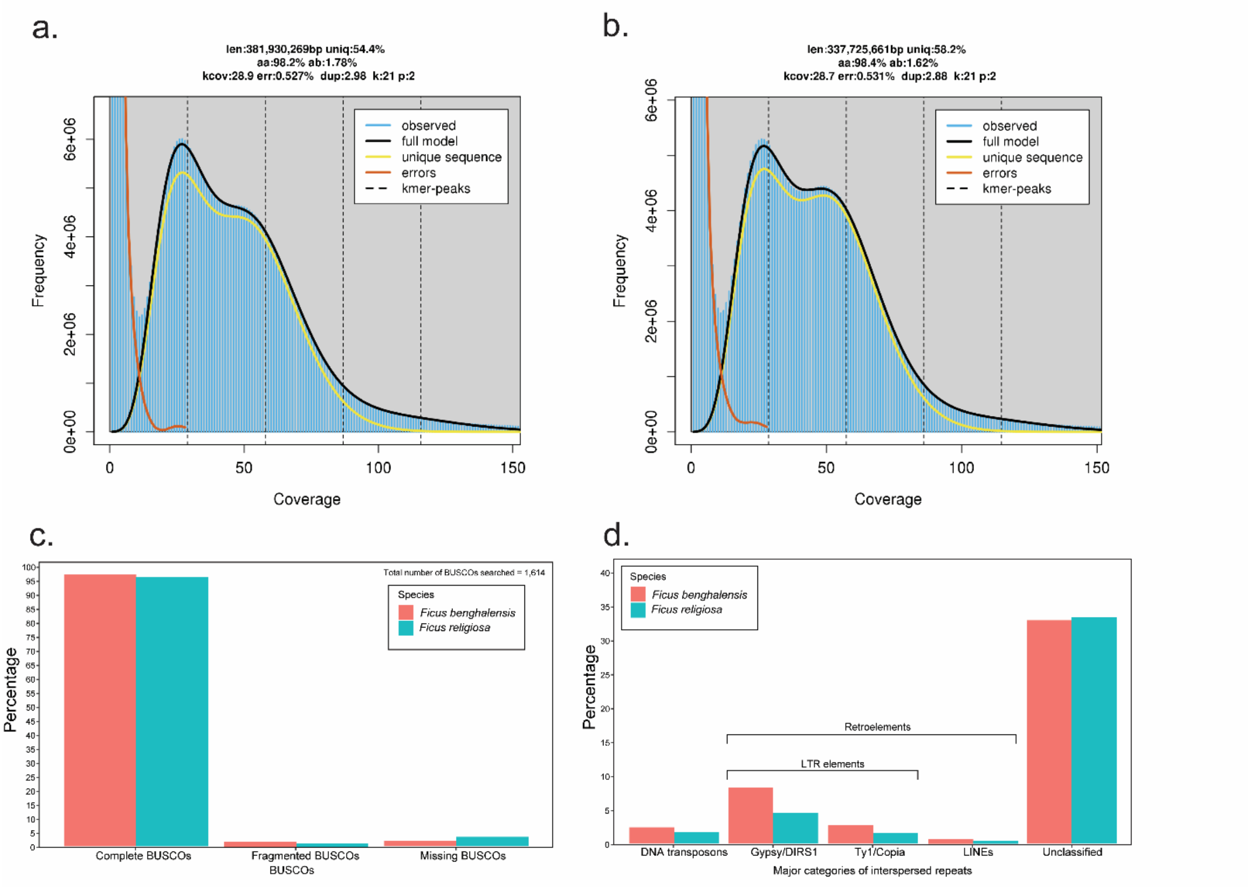

**Supplementary Figure 1.** Comparative genomic characterization of the two *Ficus* species**.** **a**. GenomeScope$^2$ profile of *F. benghalensis*, **b**. GenomeScope$^2$ profile of *F. religiosa*, **c**. BUSCO statistics of the two *Ficus* species, **d**. Repeat contents of the two *Ficus* species.

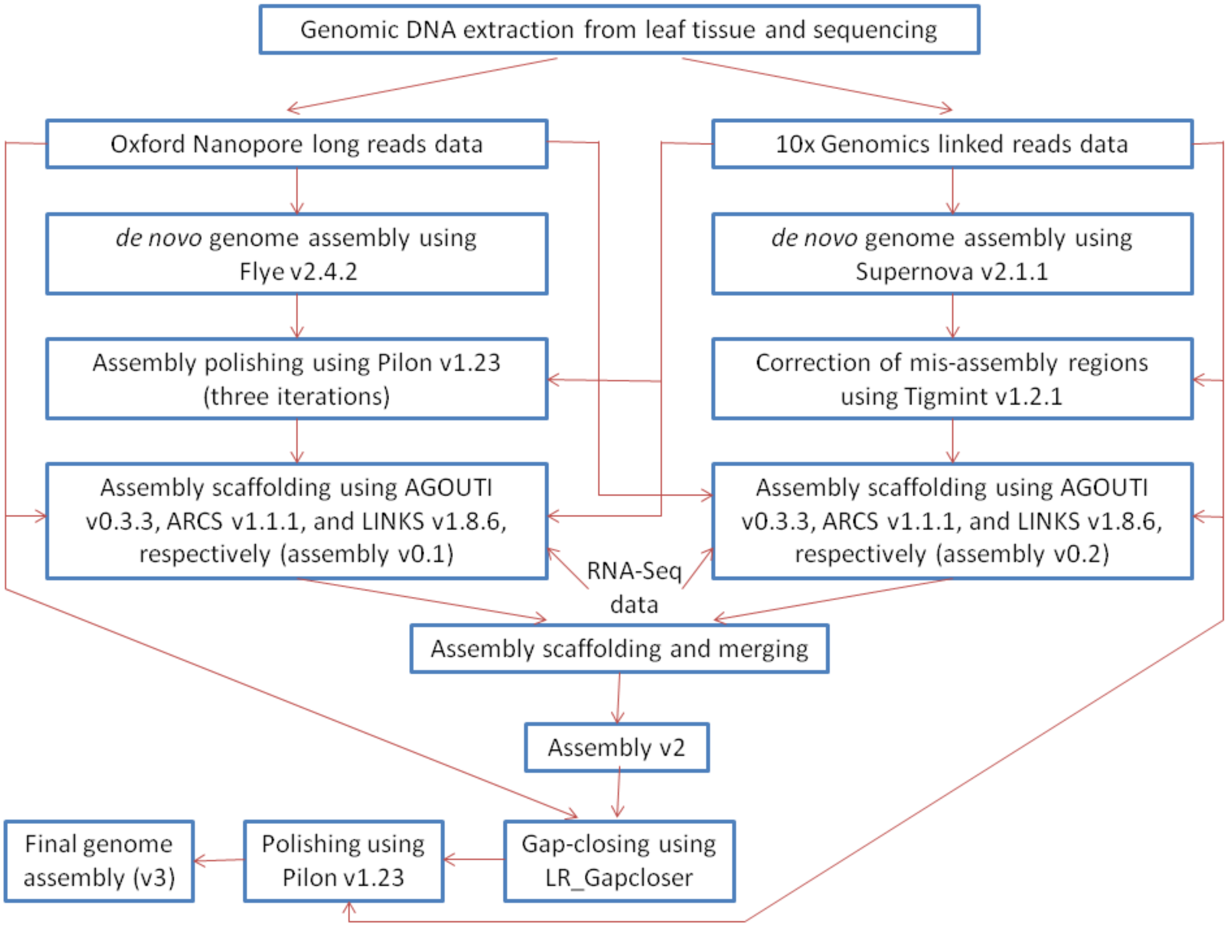

**Supplementary Figure 2.** The complete genome assembly workflow for the two *Ficus* species

**SUPPLEMENTARY NOTES**

**Supplementary Notes 1.**

**Sample Collection and DNA Extraction**

The plant samples were collected from Bhopal, Madhya Pradesh, India (23.2280252°N 77.2088987°E). The collected leaves were taken to the laboratory immediately and the leaves were cleaned before processing for DNA and RNA extraction. A few leaves were processed directly and remaining was kept frozen for later use. The leaves were homogenized in liquid nitrogen using a mortar pestle. The powdered leaves of *F. religiosa* were additionally processed by washing once with 70% ethanol followed by washing twice with autoclaved distilled water[3]. The pre-heated (at 60˚C) Carlson buffer [100mM Tris-HCl, 2% CTAB, 1.4 M NaCl, 1% PEG 8000 and 20mM EDTA (pH 9.5)] supplemented with β-mercaptoethanol (0.25 %) was added to powdered leaves that was further aided with 0.2% RNase A (20mg/mL) and 2.5% Proteinase K (20mg/mL) for removal of RNA and proteins, respectively. The tubes were then incubated at 60˚C for two hours with inverting the tubes for mixing at every 15 mins. The tubes were allowed to cool at room temperature (RT) and Chloroform (1 mL) was added, mixed by inverting the tubes and centrifuged at 5,000 xg for 15 mins at 4˚C. The supernatant was collected and 0.7X ice-cold Isopropanol was added to facilitate DNA precipitation. The precipitation was eased by an overnight incubation at -20˚C. The precipitated DNA was pelleted down by centrifuging at 5,000 xg for 45 mins at 4˚C. The DNA pellet was allowed to dissolve in G2 buffer (500 µL) of DNeasy Blood and Cell Culture kit (Qiagen, CA, USA). The dissolution was facilitated by incubation at 50˚C. The DNA dissolved G2 buffer was poured into equilibrated Genomic tip 20. Multiple tubes were added to one Genomic tip 20 in order to increase the DNA concentration. The genomic tip is then washed thrice with QC buffer (1 mL). The DNA was eluted in 1 ml of pre-heated QF buffer (pre-heated at 55˚C). 700 µL of Isopropanol was added and homogenized by inverting gently. For DNA precipitation, an overnight incubation at -20˚C was provided. The next day DNA was pelleted down by centrifuging at 5,000 xg for 15 mins at 4˚C. The DNA pellet was washed with 70% ice-cold ethanol and was allowed to air-dry the pellet. The DNA pellet was resuspended to 100 µl Nuclease Free Water (NFW). All the steps of DNA extraction were performed as gentle as possible to retain larger fragments. The quality and quantity of extracted DNA was assessed on Nanodrop 8000 spectrometer (Thermo Scientific) and Qubit 2.0 fluorometer (Invitrogen, Thermofisher Scientific, USA). The quantification was done using Qubit ds DNA BR assay kit (Invitrogen, Thermofisher Scientific, USA).

**Species identification**

For species identification, one nuclear gene i.e., Internal Transcribed Spacer (ITS) and one plastid gene i.e., Maturase K (MatK) were used. The extracted DNA was amplified for these two gene using the following primers:

1. ITS1P-F 5'-TCCGTAGGTGAACCTGCGG-3'
2. ITS2P-R 5'-TCCTCCGCTTATTGATATGC-3'
3. MatK-F 5'-CGATCTATTCATTCAATATTTC-3'
4. MatK-R 5'-TCTAGCACACGAAAGTCGAAGT-3'

The PCR programme for ITS amplification, ran on Veriti 96 well thermal cycler (Applied Biosystems, Thermofisher Scientific, USA), included initially denaturation at 94 ˚C for 3 mins, 35 cycles [ 94 ˚C (1 min), 55 ˚C (1 min) and 72 ˚C (2.5 mins) and followed by an extension at 72 ˚C of 10 mins. PCR programme for MatK ran on the same thermal cycler - initial denaturation at 95 ˚C for 3 mins, 35 cycles of denaturation at 95 ˚C for 30 sec, annealing at 50 ˚C for 3 mins and extension at 72 ˚C for 1:15 min and final extension for 7 mins at 72 ˚C. Paq5000 polymerase (Agilent Technologies, Santa Clara, CA) and Taq Polymerase (Invitrogen, Thermofisher Scientific, USA) were used for ITS gene and MatK gene amplification, respectively. The amplified products were assessed by running them on 2% agarose gel electrophoresis. The amplified products were purified and sequenced on in-house ABI Sanger sequencer at IISER Bhopal. The obtained sequences were searched for best alignment using BLASTN: Nucleotide BLAST (Basic local Alignment Search Tool) on National Center for Biotechnology Information (NCBI).

**Genomic Sequencing**

The 1.5 µg of DNA was taken for library preparation on Chromium Controller instrument using Chromium Genome Library kit with Gel Bead Kit v2 (10X Genomics, CA). The library size was assessed on Agilent Tapestation using High Sensitivity D1000 ScreenTape (Agilent Technolgies, Santa Clara, CA). The library was run on Illumina Novaseq600 sequencer (Illumina Inc., USA) for obtaining 150 bp paired-end reads.

For Nanopore Sequencing, the Genomic tip 20 (Qiagen, CA, USA)-purified DNA was further purified with Ampure XP magnetic beads (Beckman Coulter, Brea, CA). Among the purified DNA samples, only the samples obtained with the Oxford Nanopore recommended DNA quality was used for sequencing. The 1.5 µg of purified DNA was taken for library preparation using SQK-LSK109 library preparation kit

(Oxford Nanopore Technologies, UK) following the Genomic DNA by Ligation (SQK-LSK109) protocol. The library was loaded on FLO-MIN106 flowcell and sequenced on MinION sequencer using MinKNOW software (ONT, UK). The basecalling function of MinKNOW was turned off while sequencing and basecalling was performed later on using GUPPY basecaller (ONT, UK).

**Transcriptome extraction and sequencing**

For RNA extraction, the powdered leaves were taken in tubes to which 1 mL of TRIzol reagent (Invitrogen, Thermofisher Scientific, USA) was added and mixed by vortexing for 5 mins. The tubes were then allowed to stand steady at room temperature (RT) for 5 mins in order to facilitate complete disintegration of nucleoprotein complexes. After that 200 µL of chloroform was added, vortexed for 15 sec and incubated at RT for 10 mins. The tubes were centrifuged at 12,000xg for 15 min at 4˚C to obtain the supernatant.  The supernatant that contained RNA was collected in a new tube and was allowed to precipitate by adding 500 µL of ice-cold isopropanol. The RNA precipitation was facilitated by mixing and incubating at RT for 5-10 mins. The RNA was pelleted down by centrifuging at 12,000xg for 10 mins at 4˚C. The RNA pellet was washed with 75% ethanol and residual ethanol was allowed to evaporate via air-drying. Finally, the RNA pellet was resuspended in 50 µL of NFW and the pellet dissolution was ensured by incubating at 55-60˚C for 10-15 mins. The extracted RNA was quantified on Qubit 2.0 Fluorometer using Qubit RNA HS assay kit (Invitrogen, Thermofisher Scientific, USA). The RNA library was prepared using TruSeq Stranded Total RNA library preparation kit (Illumina, CA, USA) with Ribo-Zero workflow and library size was assessed on Agilent Tapestation using High Sensitivity D1000 Screentape (Agilent Technologies, Santa Clara, CA). The library was sequenced on Novaseq 6000 platform (Illumina Inc., USA) for generating 150 bp paired-end reads.

**Supplementary Notes 2.**

**Transcriptome data pre-processing**

Paired-end transcriptome reads for the two *Ficus* species generated from Illumina sequencing platform in this study, and other previous study[4] (for *F. religiosa* species) were quality-filtered using Trimmomatic[5] v0.38 with the following parameters – "LEADING:20 TRAILING:20 SLIDINGWINDOW:4:20". Also, the reads shorter than 60 bases were discarded.

**Genome assembly**

Prior to genome assembly of the two *Ficus* species, genome size estimation was performed for both the species using 10x Genomics linked reads. Barcode sequences were filtered out using two python scripts process_10xReads.py and filter_10xReads.py, available in proc10xG (https://github.com/ucdavis-bioinformatics/proc10xG). The barcode-filtered reads were used to estimate the genome sizes using SGA-preqc (with 'ropebwt' and '--no-reverse' indexing options)[1]. The barcoded reads were also processed using Longranger basic v2.2.2 (https://support.10xgenomics.com/genome-exome/software/pipelines/latest/installation), and were used for other steps in the genome assembly process.

Adapter-processed Nanopore long reads and barcoded 10x Genomics linked reads were used separately to perform initial genome assemblies. In case of Nanopore reads-based genome assembly, the best assemblies were obtained using Flye[6] v2.4.2, in terms of genome assembly statistics. The barcode-processed 10x Genomics linked reads were used to polish (three times) the assemblies derived from error-prone Nanopore reads (to fix erroneous bases, indels, mis-assemblies) using Pilon[7] v1.23. During each iteration in Pilon, barcode-processed reads were mapped to the genome assembly using BWA-MEM[8] (v0.7.17), and SAMtools[9] (v1.9) was used to process and sort these alignments. Similarly, in case of 10x Genomics reads-based assembly, Supernova[10] v2.1.1 was used to obtain the haplotype-phased assemblies (with maxreads=all, and 'pseudohap2' style). The mis-assembly regions in the Supernova-derived assemblies were corrected using Tigmint[11] v1.2.1 (a combination of tigmint-molecule and tigmint-cut) using the barcode-processed read mapping-based alignments obtained from BWA-MEM, and SAMtools.

Both the genome assemblies constructed from Nanopore and 10x Genomics reads were scaffolded using quality-filtered paired-end RNA-Seq reads, barcode-processed 10x Genomics reads, and error-corrected (using Canu[12] v2.0) Nanopore reads, respectively. RNA-Seq reads were mapped on the polished assemblies to generate the read alignments using BWA-MEM and SAMtools. Using these alignments and ".gff3" files (obtained using AUGUSTUS[13] 3.2.3), scaffolding was performed using AGOUTI[14] (v0.3.3). Further scaffolding was performed using ARCS[15] v1.1.1 with the 10x Genomics read alignments (obtained using Longranger align) on the assemblies. Canu-corrected Nanopore long reads were also used for the next round of scaffolding using LINKS[16] v1.8.6 (with default parameters), to construct assembly v0.1 (from Nanopore reads) and assembly v0.2 (from 10x Genomics reads). These two scaffolded genome assemblies were merged into a higher contiguous single assembly using Quickmerge[17] v0.3 (assembly v1), and BLASTN was also used for merging not to miss any unique contig[18]. In case of BLASTN merging, stringent parameters were used as following - sequence identity 10%, query coverage 10%, e-value $10^{-9}$ (for both the species). Contigs from assembly v0.2 that did not match to assembly v1 even with these parameters were added to assembly v1 to construct assembly v2. Gap-closing and final assembly polishing was performed on assembly v2 to construct the final genome assemblies for the two *Ficus* species (assembly v3).

**Identification of tandem repeats and non-coding RNAs**

Tandem Repeat Finder[19] (TRF) v4.09 was used to identify the simple repeats in the genome assemblies (assembly v3) of *F. benghalensis and F. religiosa* using the parameters –

Matching weight - 2, indel penalty - 7, mismatching penalty - 7, indel probability - 10%, match probability - 80%, maximum period size - 2000, minimum alignment score - 50.

For *de novo* prediction of tRNAs, tRNAscan-SE[20] v2.0.7 was used with default parameters, and the summary statistics for both the *Ficus* genomes are as follows -

| | *F. benghalensis* | *F. religiosa* |
|---|---|---|
| tRNAs decoding Standard 20 amino acids | 679 | 668 |
| Selenocysteine tRNAs (TCA) | 0 | 0 |
| Possible suppressor tRNAs (CTA, TTA, TCA) | 1 | 0 |
| tRNAs with undetermined/unknown isotypes | 5 | 6 |
| Predicted pseudogenes: | 46 | 51 |
| Total tRNAs | 731 | 725 |
| tRNAs with introns | 33 | 30 |

Hairpin miRNAs were identified in both the *Ficus* species with a homology-based approach using miRBase database[21]. A total of 22,365 hairpin miRNA sequences were clustered from a set of 38,589 hairpin miRNA sequences (available in miRBase database) using CD-HIT-EST[22] v4.8.1 (sequence identity 90%), which were used for BLASTN search (sequence identity 80%, e-value $10^{-3}$) to identify the hairpin miRNAs in the *Ficus* genomes.
